# Color naming and categorization depend on distinct functional brain networks

**DOI:** 10.1101/2020.04.13.038836

**Authors:** Katarzyna Siuda-Krzywicka, Christoph Witzel, Paolo Bartolomeo, Laurent Cohen

## Abstract

Naming a color can be understood as an act of categorization, i.e. identifying it as a member of category of colors that are referred to by the same name. But are naming and categorization equivalent cognitive processes, and consequently rely on same neural substrates? Here, we used task and resting-state fMRI, as well as behavioral measures to identify functional brain networks that modulated naming and categorization of colors. Color naming and categorization response times were modulated by different resting state connectivity networks spanning from the color-sensitive regions in the ventro-occipital cortex. Color naming correlated with the connectivity between the left posterior color region, the left medial temporal gyrus, and the left angular gyrus; whereas color categorization involved the connectivity between the bilateral posterior color regions, the left frontal, right temporal and bilateral parietal areas. The networks supporting naming and categorization did not overlap, suggesting that the two processes rely on different neural mechanisms.

**Significance:** When we name a color, we also identify it as a member of a color category, e.g. blue or yellow. Are neural processes underlying color categorization equivalent to those of color naming? Here, we address this question by measuring how individual differences in color categorization and naming response times relate to the strength of functional connections in the brain. Color naming speed correlated with left-hemispheric connectivity between the color-sensitive visual regions and the anterior temporal lobe. Color categorization speed was modulated by a different brain network, encompassing bilateral color-sensitive visual areas, and high-level executive and semantic regions. Thus, color categorization and naming performance involved distinct, non-overlapping brain networks, suggesting that the two processes depend on different neural mechanisms.

## 1. Introduction

Does the human ability to efficiently categorize visual stimuli rely on language? Color categorization, i.e. our ability to group colors into ensembles such as green, yellow and red, has served as a case-in-point of this debate (for reviews, see: Witzel, 2018; Lindsey and Brown, 2019; Siuda-Krzywicka, Boros, *et al.*, 2019). Studies showed that language regions operate when people categorize colors (Ting Siok *et al.*, 2009) and that color naming may be necessary for the visual system to respond to color categorically (Thierry *et al.*, 2009; Athanasopoulos *et al.*, 2010; Brouwer and Heeger, 2013). Yet, other studies did not find evidence for the engagement of language or language regions in color categorization (see e.g. Bird *et al.*, 2014; Persichetti *et al.*, 2015; for review see Siuda-Krzywicka *et al.*, 2019). In patients with naming deficits due to brain damage, performance in certain color categorization tasks can be significantly better than in color naming (Roberson, Davidoff and Braisby, 1999; Haslam *et al.*, 2007; Siuda-Krzywicka, Witzel, *et al.*, 2019). Particularly, a patient with color anomia due to a left ventro-occipital lesion had severely impaired color naming and close-to-normal performance in color categorization (Siuda-Krzywicka, Witzel, *et al.*, 2019). Different brain circuits may then support categorization and naming.

We assessed the hypothesis of distinct brain mechanisms by using functional connectivity and behavioral measures. Individual patterns of connectivity show pronounced inter-subject variability, specific enough to allow identifying individual subjects by their connectivity fingerprints (Finn *et al.*, 2015). Moreover, for a wide variety of cognitive functions, correlations exist between individual behavioral scores and the connectivity of the involved brain regions (see Thiebaut de Schotten *et al.*, 2011 for visuo-spatia attention; Finn *et al.*, 2015 for fluid intelligence; Galeano Weber *et al.*, 2017 for working memory; Stevens *et al.*, 2017 and López-Barroso *et al.*, 2020 for reading efficiency; Rosenberg *et al.*, 2020 for sustained attention). The strength of brain connectivity may be sensitive to even small behavioral differences, such as the amount of words read per minute (López-Barroso *et al.*, 2020).

Using a similar approach, we used resting state functional connectivity to study whether response time variability in color naming and color categorization is linked to neural variability in identical or in distinct brain systems. To this end, we first identified occipitotemporal color-sensitive regions, and assessed individual performance in a non-verbal color categorization task and a verbal color naming task. We then studied the functional connectivity of those color regions, focusing on the left color regions, linked to color naming in lesion studies (Siuda-Krzywicka and Bartolomeo, 2019). We showed that distinct, non-overlapping resting-state networks are correlated with response times in color naming and in color categorization. Moreover, we showed that this dissociation fits with the brain lesions and the connectivity pattern observed in a patient with color anomia but spared color categorization (Siuda-Krzywicka and Bartolomeo, 2019; Siuda-Krzywicka, Witzel, *et al.*, 2019). Our results support the hypothesis that color naming and categorization are supported by distinct neural circuits in the human brain.

## 2. Results

### 2.1. Color categorization and color naming behavior

Twenty participants (6 females, aged 49.5±7.24 years, see Methods 4.1. for details) took part in the study. 18 of them performed a behavioral color categorization task and a color naming task (Fig. 1A, Siuda-Krzywicka, Witzel, *et al.*, 2019). In the categorization task, two vertically arranged, bipartite discs were presented on each trial. Each disk contained two colors, either from the same color category (e.g., two shades of blue) or from different categories (e.g., brown and red). Participants had to indicate the disc containing the same-category colors. The same displays were used for the color naming task. Participants heard a color name and had to indicate the disc containing the corresponding color. The colors used in the study were classified as a member of a given category with above 90% consensus in a separate color naming experiment (for details, see Methods 4.2. and Siuda-Krzywicka, Witzel, *et al.*, 2019). Overall, in the color categorization task, the participants were 89±4% accurate and on average responded in 2.08±0.64 seconds. In the color-naming task, they responded with 99±1% accuracy with an average response time of 1.12±0.2s. There was virtually no variance in color naming accuracy (σ^2^= 0.0002), preventing statistical inference on this variable, as well as comparisons with color categorization accuracy.

**Figure 1.**
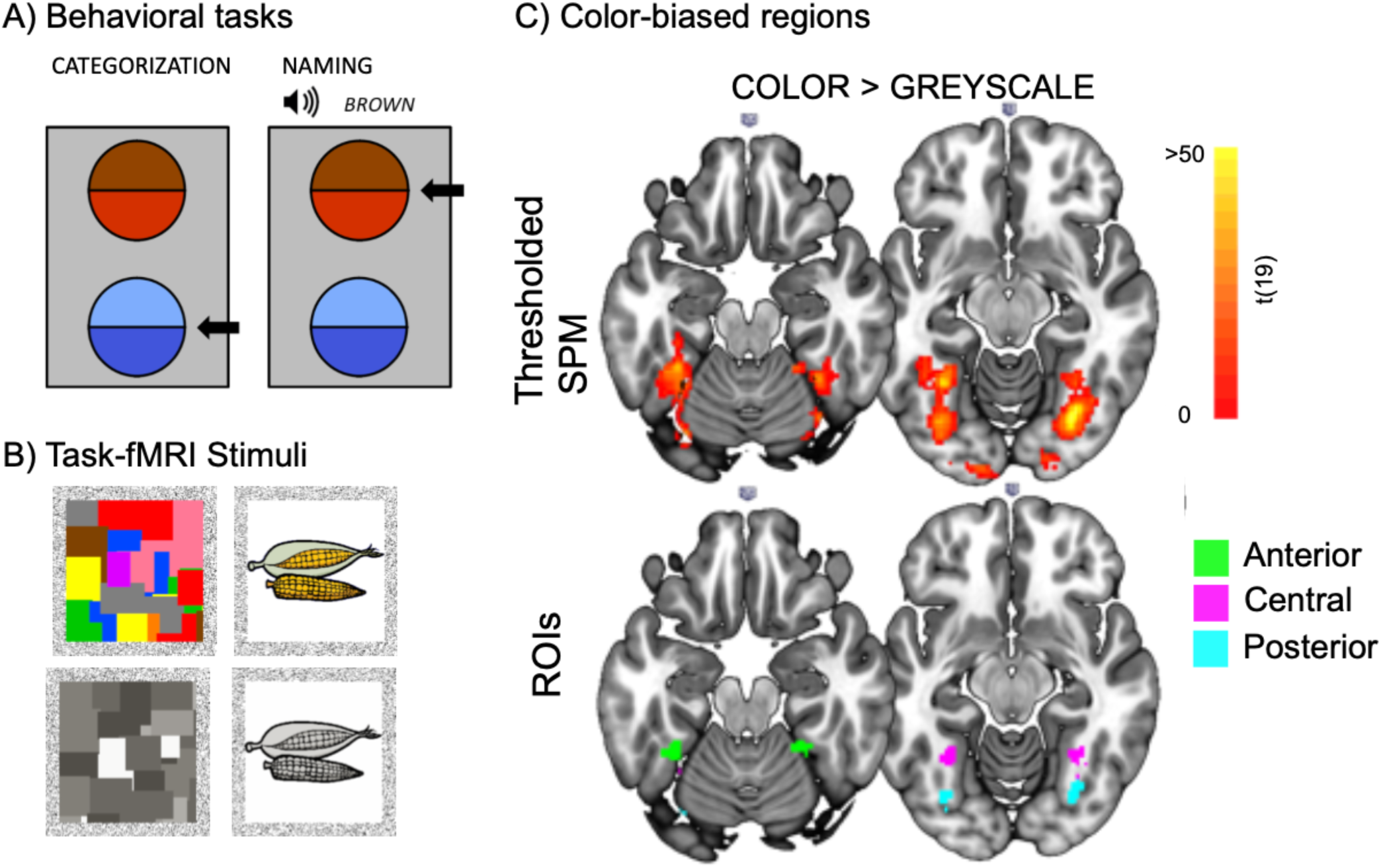
Behavioral assessment and task fMRI. (A) Example displays in the color categorization and color naming tasks. Arrows indicate correct responses. (B) Example stimuli in the task fMRI localizer experiment. (C) Color-sensitive regions defined with a contrast of color vs. greyscale. Top row: SPM map thresholded at p=0.001 voxel-wise and p=0.05 cluster-wise (with 3DClustSim correction). Bottom row: regions of interests (ROIs).

### 2.2. Color-sensitive regions in the brain localized with task-fMRI

All participants participated in a task fMRI experiment in which they viewed images of colored and greyscale objects and abstract Mondrians (Fig. 1B and Methods). A group-level contrast of colored minus grey-scale images revealed a medial ventro-occipital cluster of activity ranging from the early visual areas to the parahippocampal cortex. Color signals in the early visual cortices do not correspond to the perceptual color space in primates (Brouwer and Heeger, 2009; Bohon *et al.*, 2016), therefore for the subsequent analysis we focused on the areas anterior to the early visual cortices (anterior to MNI y =-80). Congruent with previous reports (Lafer-Sousa, Conway and Kanwisher, 2016), we identified three color-sensitive regions in both hemispheres, namely: a posterior color region (MNI 30 −72 −10 and −32 −76 −16), corresponding to areas V4/VO1; a central color region (MNI −32 −56 −12 and 34 −50 −20), adjacent to the anterior border of area VO2; and an anterior color region (MNI −34 −44 −22 and 22 −44 −20) at the border of the anterior fusiform and parahippocampal gyri (Fig. 1C). This allowed us to define six regions that served as seeds for the subsequent connectivity analyses. Each seed consisted in the 50 voxels with the highest activation (beta value) located within 8 mm of the above peaks (see Fig. 1C and Methods 4.3).

### 2.3. Resting state connectivity of the color-sensitive regions

We first investigated the overall functional connectivity of the 6 color-sensitive seeds defined before, using a whole brain omnibus F test (Fig. 1C, see Methods 4.4). Connected regions encompassed the entire occipital cortex, including both ventral and dorsal aspects; and more distant brain areas in the bilateral superior temporal gyri and frontal cortex (Fig. 2A; p=0.001 voxel-wise, p=0.05 FDR corrected cluster-wise).

**Figure 2.**
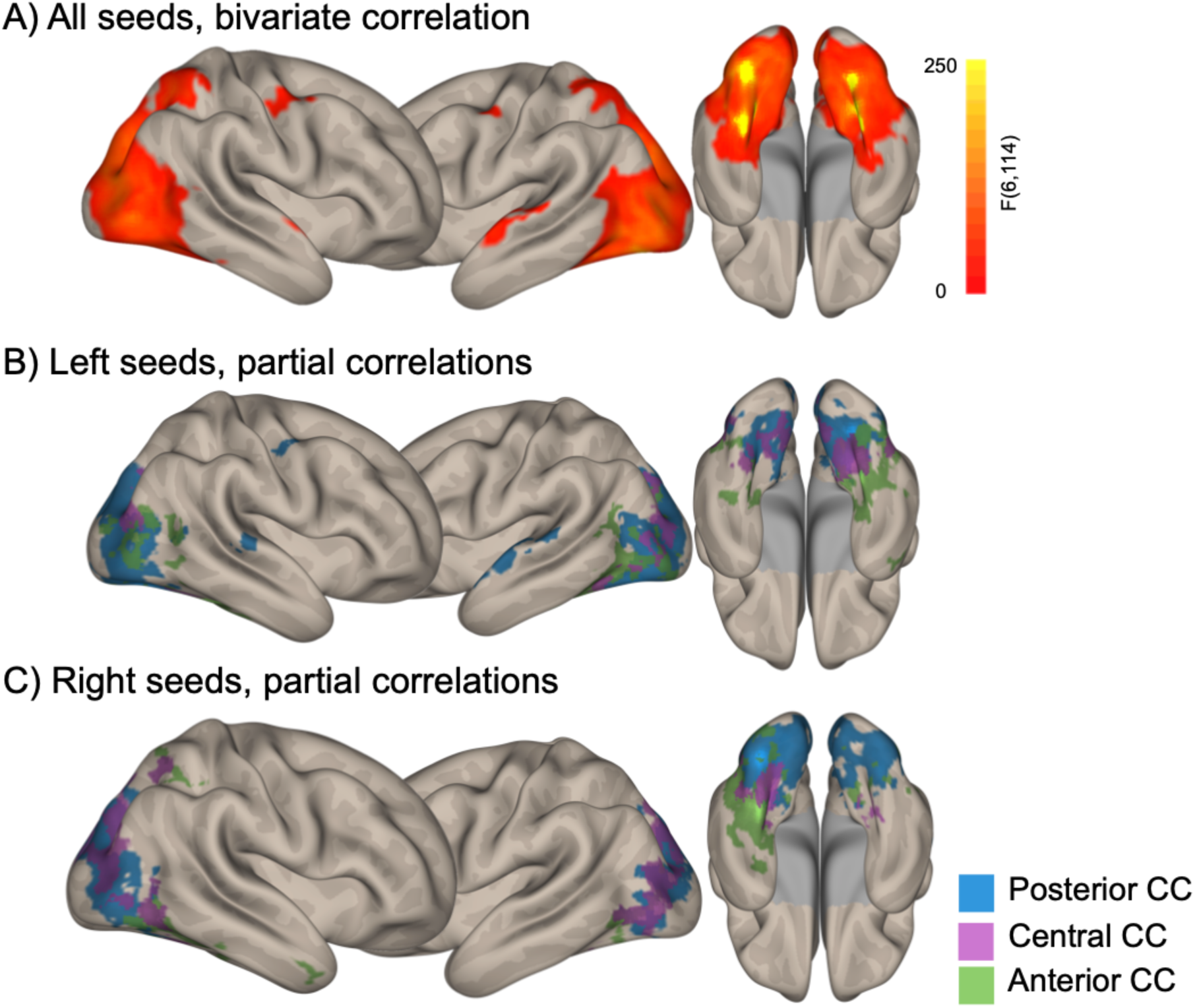
Resting-state connectivity of the six ROIs. (A) Brain regions connected to at least one of the six color seeds, as computed with an F test. (B-C) Unique connectivity patterns as assessed with partial correlations, for the left- (B) and right-hemispheric (C) seeds. Voxelwise p<0.001, cluster wise p<0.05 FDR corrected.

To characterize the unique connectivity pattern of each seed, we used partial correlations. For each seed, we identified connected regions while partialling out the contribution of all other seeds (see Methods 4.4). This analysis revealed a significant and unique pattern for each color seed (Fig. 2 B,C; p=0.001 voxel-wise, p=0.05 FDR corrected cluster-wise), beyond its immediate surrounding. Particularly, the left posterior color seed was connected to the left superior temporal gyrus (Fig 2B), while the anterior color seeds in both hemispheres were connected to the ventral temporal lobes and the parahippocampal cortex.

### 2.4. Color naming and functional connectivity of the left-hemispheric color-sensitive regions

To investigated how individual differences in color naming response times (RTs) relates to the connectivity pattern of left-hemispheric seeds, we set up a linear regression model. In this model, voxel-wise connectivity values with left color-seeds served as dependent variable, and participants’ age, gender and mean log-transformed RTs in the color naming task as predictors (see Fig. 1A and section 2.1). For clarity, we converted regression beta coefficients for color naming RTs to partial correlation coefficients, controlling for the effects of age and gender (see Methods 4.4)

Figure 3A shows the correlation maps of the left color seeds, i.e. regions whose connectivity with the left color seeds was correlated with color naming RTs (see Methods 4.4 for details). Shorter color naming RTs corresponded to stronger connectivity between the left color seeds and the left-predominant middle temporal gyrus, the left angular gyrus, the left precentral and supramarginal gyri (p=0.01 voxel-wise, p=0.05 cluster-wise; for right-sided color seeds see fig 4C). To investigate the specific effect of each of the left color seeds, we performed t tests comparing the correlation maps of one color seed to the average correlation map of the remaining two. The left posterior seed connectivity was more correlated with color naming RTs in the left middle temporal gyrus, the left angular gyrus and anterior frontal regions (Fig. 3B and S1B, p=0.01 voxel-wise and 0.05 cluster-wise FDR corrected; for the corresponding map of the left anterior seed, see Fig S1C). These left middle temporal clusters overlapped with the regions that were functionally disconnected from color-sensitive visual cortex in a patient with color anomia (see Fig. 3C and Siuda-Krzywicka, Witzel, *et al.*, 2019). Accordingly, participants who were faster at naming colors had stronger functional connectivity between the left posterior seed and the anterior temporal clusters disconnected in this patient (Fig 3 C-D, r(14)= −0.65, p=0.009, see Methods 4.4).

**Figure 3.**
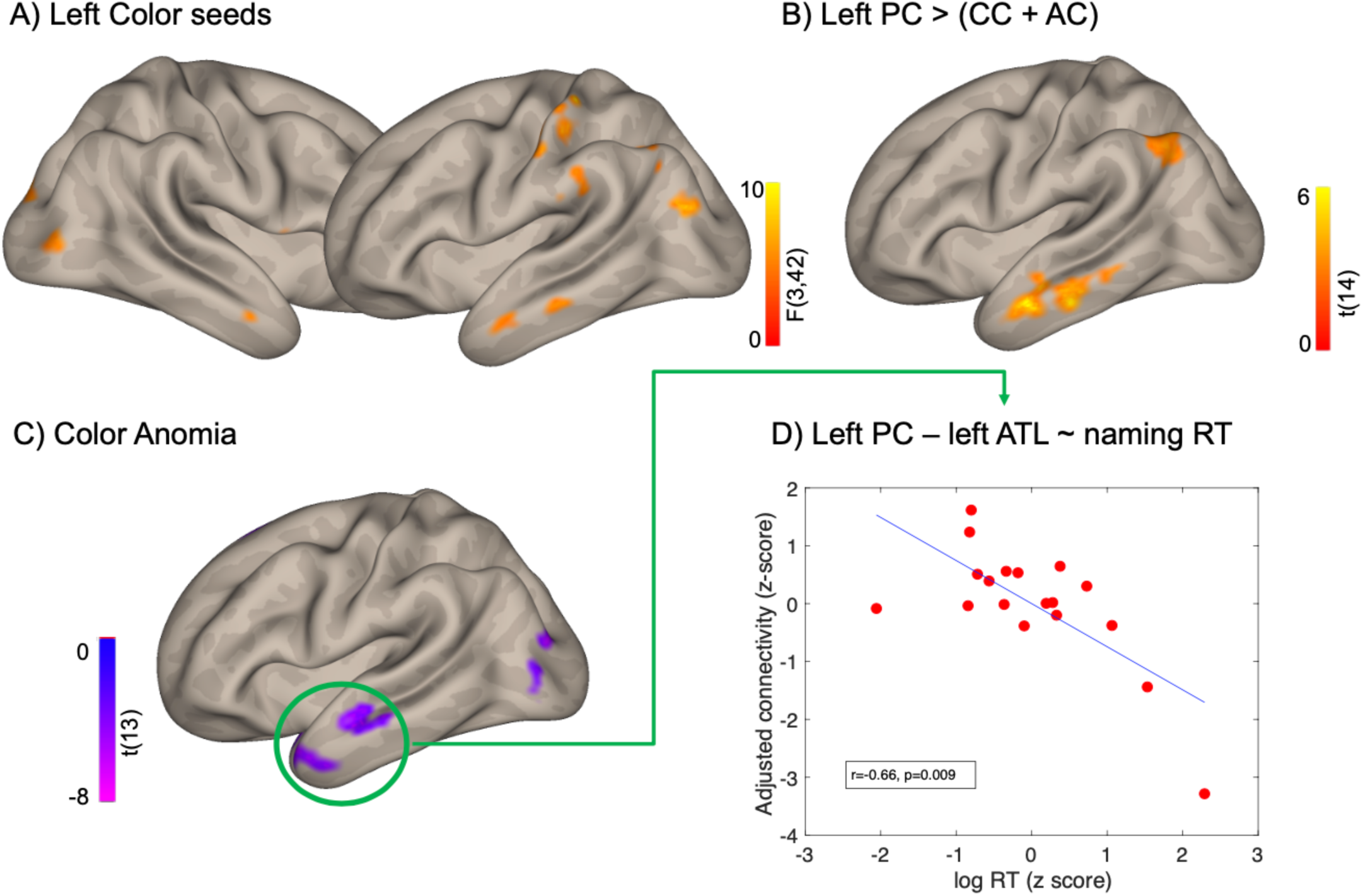
Correlation between functional connectivity and color naming response times. (A) Areas where the connectivity to at least one of the left color-seeds is modulated by color naming response times (F test, see Methods 4.4). (B) Brain areas where correlation with naming RT is stronger for the left posterior seed than for the anterior and central color seeds. (C) Brain regions significantly disconnected from color-sensitive cortex in a color anomic patient (green circle, Siuda-Krzywicka, Witzel, *et al.*, 2019). (D) Correlation between naming RT and the connectivity (adjusted for participants’ age and gender) of the left posterior color seed with the disconnected regions highlighted in (C). Voxel-wise thresholds: (A, B) 0.01 (C) 0.005; cluster-wise thresholds (A) p<0.05 uncorrected; (B,C) p<0.05 FDR corrected.

**Figure 4.**
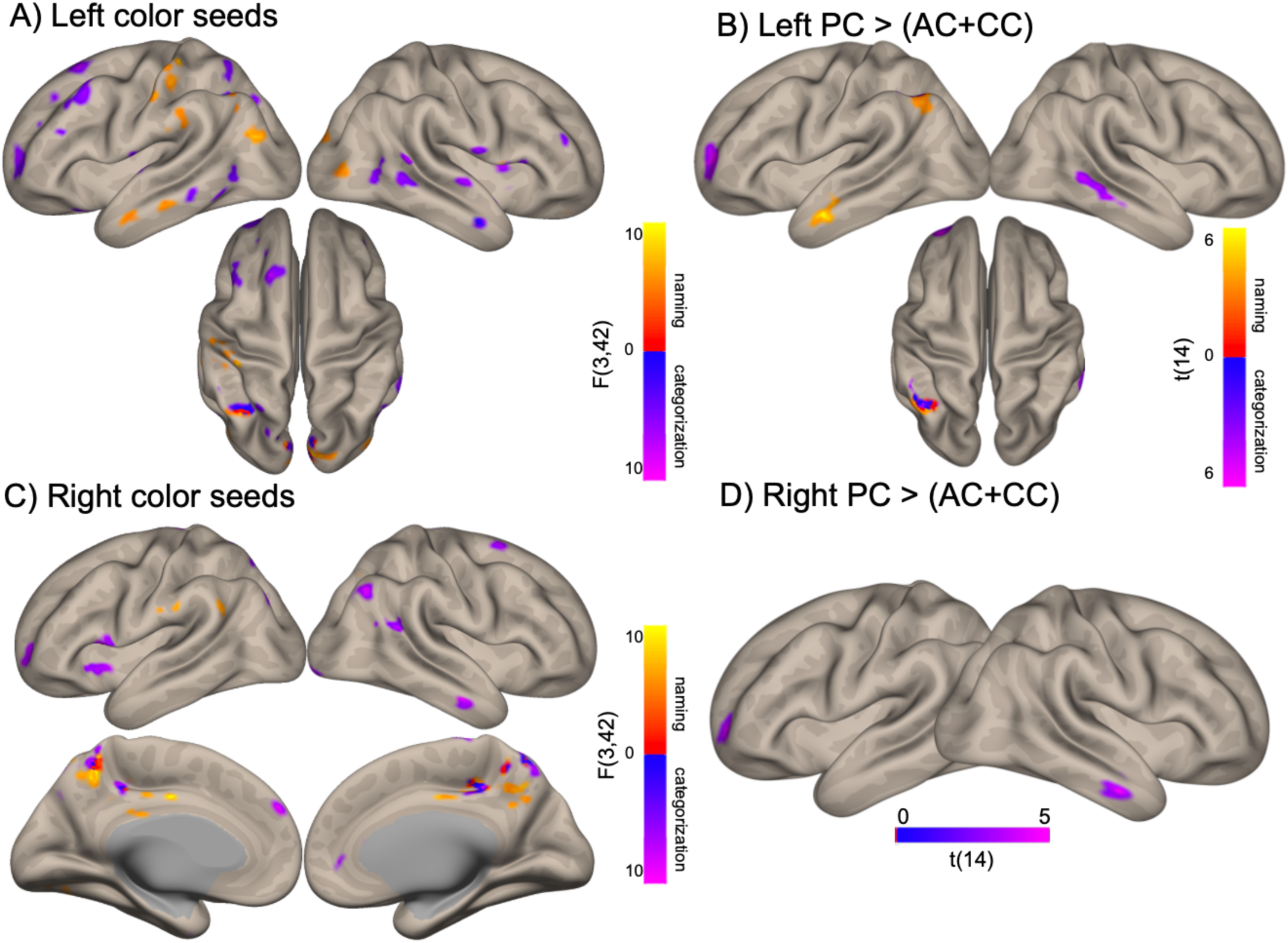
Distinct networks for color categorization and color naming. (A, C) Correlation of the connectivity of the left (A) and right (C) color seeds with RTs in categorization (blue-purple) and naming (red-yellow) tasks (B, D) Regions where the effects of categorization and naming RT were greater for posterior seed than the anterior and central seeds, in the left (B) and right (D) hemisphere. Voxel-wise threshold (A,C) p=0.01; (B,D) p=0.005; Cluster-wise thresholds (A,C) p=0.05 uncorrected; (B,D) p=0.05 FDR corrected.

Analyses using an anatomically-defined Regions of Interest (ROI) confirmed the above voxelwise results (Fig. 3D and Methods 4.4): Shorter color naming RTs corresponded to stronger connectivity between the left posterior seed and the left middle temporal gyrus (r(14)= −0.57, p=0.026 for its anterior portion and r(14)= −0.52, p=0.038 for its posterior portion).

We then investigated whether the link between naming RTs and functional connectivity was specific to seeds located in color-sensitive cortex. To this end, we computed the correlation of naming RTs with the connectivity of the left middle temporal gyrus with the left Lateral Occipital Complex (LOC, MNI −50 −68 −6) and with the Visual Word Form Area (VWFA, MNI −44 −40 −20, see Fig. S1C). Neither of those correlations were significant (see Table S1). This supports a specific involvement in color naming of the coupling between the left posterior color seed and the left middle temporal gyrus.

### 2.5. Color categorization and functional connectivity of the left color-sensitive regions

Following the same method as in the previous section, we studied the correlation between individual RTs from the color categorization task (Fig 1A) with the functional connectivity of color seeds. To compare resting state networks involved in color naming to the networks involved in color categorization, we focused on the left color seeds, and the posterior color seed in particular. Figure 4 A-B presents the overlap between categorization and naming correlation maps involving the left seeds. Shorter categorization RTs corresponded to stronger connectivity between the left color seeds and the left superior frontal gyrus, the left superior parietal cortex, bilateral inferior frontal and temporal areas as well as the occipital pole (Fig. 4A, medial occipital connectivity non-visible in the figure, p=0.01 voxelwise; p=0.05 cluster-wise). Compared to the other two left color seeds, the posterior seed’s connectivity was more correlated with RTs in the left anterior middle frontal gyrus and the right middle temporal gyrus (Fig. 4B, for the corresponding map of the left anterior seed, see Fig S1D).

Importantly, the networks identified for the naming and categorization tasks were very different. First, regions where left seeds connectivity correlated with categorization RTs were distributed across both hemispheres, whereas regions correlated with naming RTs predominated in the left hemisphere (Fig 4 A, B). Second, there was a minimal overlap between regions involved in the two tasks, even when thresholding maps at p=0.01 voxel-wise (Fig 4 A).

### 2.6. Exploratory analysis of color naming and categorization effects on functional connectivity of right color seeds

To explore possible overlaps between categorization and naming brain networks stemming from the right color seeds, we repeated the same analyses, but using the connectivity patterns of right seeds as the dependent variable. In Fig 4C we plotted the two correlation maps and their overlap (both p=0.01 voxel-wise and p=0.05 cluster-wise). Color categorization RTs correlated with right sided color cortex connectivity to distributed frontal regions (left inferior and anterior middle frontal gyri, right superior frontal gyrus), right supramarginal, angular and middle temporal gyri (Fig 4C). Regions where right color seeds connectivity correlated with color naming RTs were limited to the medial parietal/posterior cingulate cortex and left parietal operculum (Fig 4C). Interestingly, left and right posterior seeds showed stronger color categorization RTs modulation of connectivity with similar left frontal and right temporal regions (see purple clusters in Fig 4B and D, p<0.005 voxel-wise, p<0.05 FDR-corrected cluster-wise).

## 3. Discussion

We investigated the functional connectivity patterns of the three color-sensitive regions lying in the bilateral ventral occipito-temporal cortex, namely the posterior, central, and anterior color regions. We showed that those regions have distinct connectivity patterns that encompass both early visual cortices and high-level supramodal frontal and temporal areas. These distinct connectivity patterns indicate that the three color regions may have different functional profiles, an idea put forward in the literature of high-level color processing (Conway, 2018; Siuda-Krzywicka and Bartolomeo, 2019). Here, we show that the connectivity of the color regions shows different spatial patterns of correlation with response times in naming and categorization tasks.

### 3.1. Color categorization without language

To our knowledge, we present here a first attempt at identifying the neural substrate of color naming and categorization using brain connectivity measures. We showed, on the one hand, that color naming speed was related to the connectivity between the left middle temporal and angular gyri and a very specific sector of the visual cortex, namely the left posterior color region. On the other hand, color categorization speed was related to the connectivity between left frontal, right temporal, bilateral parietal areas, and the bilateral posterior color regions. Consistent with our previous report (Siuda-Krzywicka, Witzel, *et al.*, 2019), all the structures involved in categorization (apart from the left posterior color region) were spared in a severely color anomic patient with spared color categorization. Overall, we demonstrate that different neural systems underlie naming and categorization, and that the two processes can show dissociated impairments (see also Garcin, Volle, *et al.*, 2018 for categorization deficits in frontal patients without naming impairments).

Color categories could rely on domain-general mechanisms of semantic knowledge organization, rather than being a purely perceptual or linguistic process (reviewed in: Siuda-Krzywicka, Boros, *et al.*, 2019). The frontal portions of color categorization networks identified in this study agree with findings in humans and macaques (Koida and Komatsu, 2007; Bird *et al.*, 2014), and suggest that color categorization may rely on brain systems engaged in attention and task-demands. Attention could also explain the category effect on color discrimination, i.e. easier discrimination of equidistant colors stemming from different categories, as opposed to same category colors (Witzel and Gegenfurtner, 2015). The right temporal component of the categorization network reported here is congruent with a recent finding that cortical thickness of the right anterior temporal lobe was modulated by individual differences in response times in a non-color categorization task (Garcin, Urbanski, *et al.*, 2018). Based on the above, we would like to propose that color categorization can be viewed in the framework of controlled semantic cognition (Lambon Ralph *et al.*, 2017). In this framework, semantic cognition depends on the interaction between executive processes subtended by frontal regions, and semantic content resulting from the interplay between anterior temporal semantic “hubs”, and sensory “spokes” providing perceptual inputs. Posterior color regions could play the role of spokes, and provide perceptual color input to the right temporal cortex holding the content of color categories. Fronto-parietal regions would play a crucial role in accessing, retrieving and executively manipulating color categorization through attention allocation and adaptation to task demands. This model does not entail a contribution of language, and could therefore explain categorical responses to color in both humans and non-human primates, and consequently provide a universal, language-independent neural framework for color categorization (Siuda-Krzywicka, Boros, *et al.*, 2019).

### 3.2. A color naming hub ?

We showed that color naming speed correlates with the connectivity between the left posterior Color region and the left middle temporal gyrus. This result prevailed only for the left posterior color region, and we found no such correlation in the neighboring Lateral Occipital Complex or the Visual Word Form Area (see Fig S1A and table S1). The middle temporal gyrus cluster revealed in our analyses overlapped with regions disconnected from color-sensitive visual cortex in a patient with color anomia (Siuda-Krzywicka, Witzel, *et al.*, 2019), and the participants’ color naming speed correlated strongly with the connectivity of the posterior Color region and this disconnected region (see Fig 3C-D). These converging results indicate that the communication between the left posterior color region and the left middle temporal gyrus is causally related to color naming abilities. We suggested that the left sided color-sensitive regions may act as a cortical naming hub linking color perception with color names (Siuda-Krzywicka and Bartolomeo, 2019; Siuda-Krzywicka, Witzel, *et al.*, 2019). The current results allow us to narrow down this hypothesis to the posterior color region, which would serve as the color naming hub, and whose lesions would lead to color anomia.

Note that the left posterior color region was not connected with the left middle temporal gyrus when averaging all subjects (without accounting for the color naming speed, see Fig 2). This is due to the large variability in connectivity strength between our participants. As visible in Figure 3D, individual connectivity ranged from positive to negative values, yielding no significantly positive connectivity for the group as a whole. According to our results, the strength of this connectivity modulates naming speed. Hence, we predict that color naming experts, who are trained by their profession to identify and communicate colors by names, should show higher connectivity between the left posterior color region and left middle temporal gyrus.

### 3.3. Using Response Times as a measure of individual variability in color naming and color categorization

We used response times in color naming and categorization as a proxy for participants’ color categorization and color naming ability. This index may also be sensitive to individual motivation, visuo-motor processing speed or domain-general language processing speed. However, those parameters are unlikely to account for our results. Indeed, we limited the analyses to networks seeded from color-sensitive regions, ensuring that effects were specific to cortical color processing. Also supporting the specificity of our findings, we found no correlation between color naming RTs and the connectivity of neighboring occipital areas such as the VWFA or the LOC (Table S1). Moreover, we included in the analyses only the responses from the second block of the tasks (see Methods 4.2), in order to discard individual differences in how quickly participants mastered the task.

Differences in categorization and naming RTs could also reflect differences in the precise setting of the boundaries between color categories. Color categories and color names vary substantially across cultures and languages (e.g. Roberson *et al.*, 2005). Even within one culture, people vary in their choice of labels, when they describe colored items (e.g. Lindsey and Brown, 2014; Brown, Isse and Lindsey, 2016; Kuriki *et al.*, 2017), or when they sort colors into categories (reviewed in: Witzel, 2018). Response times in categorization tasks are modulated by categorical membership of colors: people are faster in categorizing and naming colors with clear category membership than colors that are more ambiguous (Bornstein and Monroe, 1980; Zimmer, 1982). In the present study, the category membership of colored stimuli was carefully controlled in an independent color naming experiment (see details in Siuda-Krzywicka, Witzel, *et al.*, 2019). As there was a full consensus on the category membership of our stimuli, it is more likely that individual differences in color naming and categorization RT reflected the speed with which colors reach the verbal lexicon and domain general categorization networks, for the two tasks respectively. This interpretation is consistent with previous reports on links between response times and brain connectivity measures (López-Barroso *et al.*, 2020), and between response times and the anatomy of brain structures relevant to categorization (Garcin, Urbanski, *et al.*, 2018).

### 3.4. Conclusions

Our findings show that color categorization and color naming are processed in separate brain networks, providing clear pathophysiological background to dissociations observed in brain-damaged patients (reviewed in: Siuda-Krzywicka and Bartolomeo, 2019). Our results suggest a novel answer to the long standing debate on the role of language vs. perception in color categorization (Siuda-Krzywicka, Boros, *et al.*, 2019). Color categorization would depend on a specific network, involving fronto-parietal executive regions, right temporal regions, and the posterior color region. This network, which goes far beyond color-sensitive regions of the visual cortex, is distinct from the color naming network, which comprises the posterior color region and the middle temporal gyrus in the left hemisphere.

## 4. Methods

### 4.1. Participants

Twenty participants (6 females, aged 49.5±7.24 years) took part in the study. All were right-handed according to the Edinburgh Inventory (Oldfield, 1971), had normal or corrected-to-normal vision, and showed normal color vision on the Ishihara Color plates test. The present research was promoted by the Inserm (CPP C13-14) and approved by the Ile-de-France I ethical committee. Before participating in this research, all participants signed an informed consent form.

### 4.2. Behavioral assessment of color categorization and color naming

The details of the color categorization and color naming procedures can be found in (Siuda-Krzywicka, Witzel, *et al.*, 2019). Each stimulus consisted in a bipartite disc, comprised of two colors (Fig 1A). The categorical membership of the two colors as well as their perceptual distance were controlled through preliminary naming and distances in CIELUV space (for details see Siuda-Krzywicka, Witzel, *et al.*, 2019). Each trial started with the presentation of a central, black fixation cross (1**°** visual angle) for 500-ms. Then two bipartite discs appeared, one above the other, aligned with the central meridian of the screen. In the color-categorization task subjects had to identify the disc containing colors from the same category. In the color-name comprehension task, the trial started with the auditory presentation of a pre-recorded color-name and the subjects’ task was to indicate the bipartite disc containing the named color. Subjects responded by pressing the upper arrow key with their right hand to indicate the upper disc, or the lower key for the lower disc. There was no time limit for responses. There were 157 trials in each experimental block. Subjects performed 2 experimental blocks of each task, but only data from the second block of each task are reported and correlated with brain connectivity measures. The response times were log transformed before correlating with connectivity measures.

### 4.3. Task fMRI

#### Behavioral procedure

For the color localizer, we used pictures from five categories (see Fig 1B): chromatic and achromatic Mondrians, and objects in congruent color (e.g. a yellow banana), incongruent color (e.g. a blue banana) and grey scale (e.g. a grey banana). For the domain localizer, we used six categories of achromatic pictures: faces, tools, houses, pairs of words, pairs of numbers and body parts. In both localizer experiments, stimuli were presented in thirty 8 second blocks alternating with 7.8 s of rest. Each stimulation block included eight pictures from one category of stimuli, displayed for 600 ms and followed by a 400 ms blank screen. Participants were asked to press a button with their right thumb whenever a picture was identical to the previous one, which was the case for 20% of stimuli (1-3 repetitions/block). For details see Siuda-Krzywicka, Witzel, *et al.*, 2019.

#### Acquisition parameters

We used a multiband echo-planar imaging sequence sensitive to brain oxygen-level-dependent (BOLD) contrast (45 contiguous axial slices, 2.5 mm isotropic voxels, in-plane matrix ¼ 80 80; TR ¼ 1022 ms; angle ¼ 62, TE ¼ 25 ms). For the color localizer, 482 volumes were acquired, for the domain localizer 576. Five additional BOLD volumes with reverse phase encoding direction were also acquired for each localizer.

#### Data preprocessing and statistical analysis

Functional images were realigned, treated with the FSL ‘‘Topup’’ toolbox in order to correct EPI distortions due to B0 field inhomogeneity (Andersson et al., 2003), normalized to the standard MNI brain space and spatially smoothed with an isotropic Gaussian filter (6 mm full width at half-maximum, FWHM).

First-level analysis was implemented in SPM12 software (https://www.fil.ion.ucl.ac.uk/spm/software/spm12/). Data were high-pass filtered and modeled by regressors obtained by convoluting the experimental conditions and the button presses with the canonical SPM hemodynamic response function (a combination of 2 gamma functions, with a rise peaking around 6 s followed by a longer undershoot). Individual contrast images were computed in order to identify (1) color-sensitive regions (colored minus grey-scale images), (2) the Visual Word Form Area (VWFA, words minus numbers), and (3) the object-sensitive Lateral Occipital Cortex (LOC, Tools minus an average of Faces and Places). Those images were smoothed with an isotropic Gaussian filter (6 mm FWHM) and entered into a second-level whole-brain one sample T-test.

### 4.4. Resting-state fMRI

#### Image acquisition

We acquired a 10-minute series of whole-brain resting-state BOLD sensitive images (gradient-echo (GE) echo planar imaging (EPI) sequence, 45 slices, slice thickness = 3mm, FOV 220 3 220mm, A>>P phase encoding direction, TR = 2990ms, TE = 26ms, flip angle = 90deg).

#### Data preprocessing

Data preprocessing and statistical analysis was performed with the CONN v.17 functional connectivity toolbox (Whitfield-Gabrieli and Nieto-Castanon, 2012). We used standard preprocessing steps including: slice-time correction, realignment, segmentation of structural data, normalization into standard stereotactic MNI space and spatial smoothing using a Gaussian kernel of 6 mm FWHM. To account for the fMRI signal attenuation, after slice-time correction, realignment, motion-correction and normalization, functional scans were subjected to intensity-based masking (Peer *et al.*, 2016; see also Siuda-Krzywicka, Witzel, *et al.*, 2019). We then used the Artifact Detection Tool (ART; https://www.nitrc.org/projects/artifact_detect/) to identify scans exceeding 3 SD in mean global intensity, and scan-to-scan motion that exceeded 0.5 mm. We regressed out those scans as nuisance covariates in the first-level analysis, together with the head motion parameters (three rotation and three translation parameters). Physiological and other spurious sources of noise were estimated with the CompCor method (Behzadi *et al.*, 2007; Whitfield-Gabrieli *et al.*, 2009; Chai *et al.*, 2012); they were then removed together with the movement- and artifact-related covariates mentioned before. We applied a temporal band-pass filter of 0.008–0.09 Hz.

The anterior and posterior MTG ROIs were defined based on the Harvard-Oxford atlas implemented in the CONN toolbox. For the anterior temporal lobe ROI identified in a patient with color anomia, a binary mask was created from a map reported in Siuda-Krzywicka, Witzel, *et al.*, 2019.

#### Statistical analysis

Seed-to-voxel whole brain connectivity first level maps were created for each participant. The average BOLD time course was extracted from six ROIs corresponding to the posterior, central and anterior color region in each hemisphere. The individual Z-maps were entered to a second-level group analysis. In the group analysis, an F test refers to a series one-dimensional contrasts, each corresponding to one seed testing against the null hypothesis that the effect of a given ROI equals to 0. To generate functional connectivity maps representing the unique connectivity patterns for each color region, we used semi-partial correlation coefficients. We calculated the connectivity values from a given seed, regressing-out the time courses of all other seeds.

We analyzed the correlations between behavior and functional connectivity by setting up a linear regression model with Fisher-r-to-z-converted connectivity values as dependent variable, in each voxel for whole-brain analyses, or averaged within a given brain region for ROI analyses. Participants’ age, gender and categorization or naming log-transformed response times were used as predictors. All predictors were z-scored. We defined separate models for naming and categorization RTs. For clarity of interpretation, we have converted the regression estimates into partial correlation coefficient using the following formula:

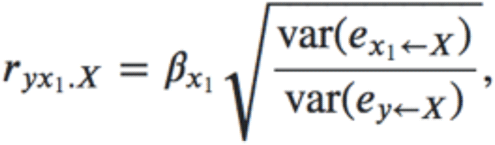

where y is the dependent variable (connectivity values), x1 is the predictor of interest (RTs), X stands for remaining predictors in the model (age and gender), B_x1_ is the regression estimate (Beta) of the RTs, and e_x1_ <- X and e_y_ <- X are the raw residuals from regressing x1 by X and y by X respectively. The partial correlation coefficients are reported within the text, the summary of regression coefficients can be found in table S1.

## Acknowledgements

This work was supported by ICM, INSERM, CNRS, the program “Investissements d’avenir” (grant number ANR-10- IAIHU-06). K.S.K. was supported by the Ecole des Neurosciences Paris Ile de France. C.W. was supported by the grant ‘Cardinal Mechanisms of Perception’ No. SFB TRR 135 from the Deutsche Forschungsgemeinschaft.

## 6. Supplementary Materials

**Figure S1.**
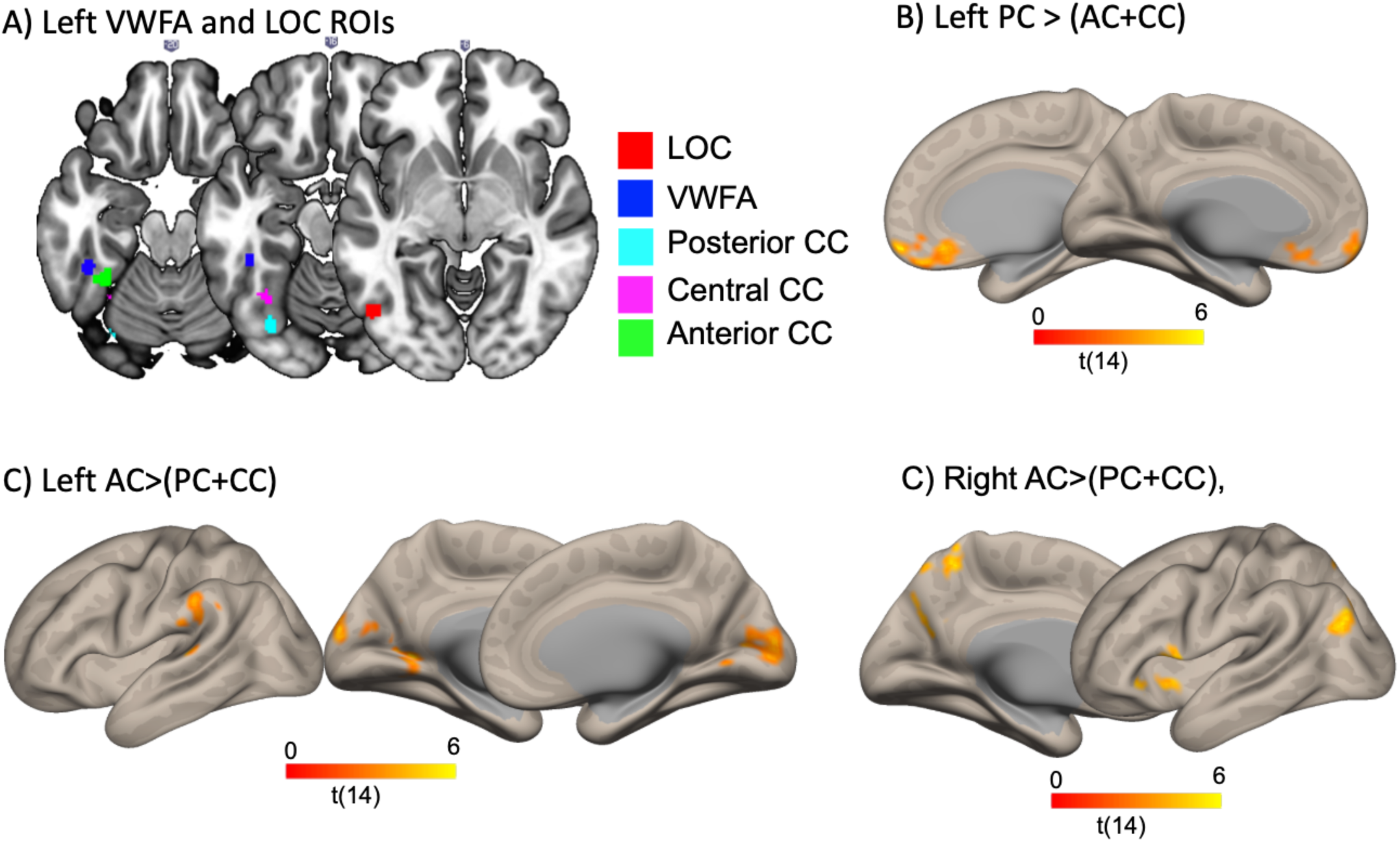
Supplementary figure associated with Figures 3 and 4. (A) The left VWFA and LOC ROIs defined from the domain localizer. The ROIs comprised 50 most activated voxels within 8mm spheres centered at peak activity for words vs. numbers for the VWFA, and tools vs. faces and places for the LOC. (B) Medial plane of Fig 3B. Note the anterior frontal clusters of connectivity. (C-D) Brain regions where RT modulation was significantly greater for the left anterior color region, compared to the other regions for color naming(C) and color categorization (D) Voxel-wise p (B-C) p=0.01, (D) p=0.005, Cluster-wise thresholds p=0.05 FDR corrected.

**Table S1 related to Fig 3.** Linear regression results of how participants’ age, gender and color naming RT explains functional connectivity between left posterior color seed (Left PC), left Lateral Occipital Complex (Left LOC) and the Visual Word Form Area (VWFA) and the left middle temporal gyrus (anterior and posterior portion as defined in the Harvard-Oxford Atlas, and the clusters of dysconnectivity in color anomia in Fig 3 C).

|  | Left PC |  |  |  | Left LOC |  |  |  | VWFA |  |  |  |
| --- | --- | --- | --- | --- | --- | --- | --- | --- | --- | --- | --- | --- |
|  | Estimate | SE | t statistics | p value | Estimate | SE | t statistics | p value | Estimate | SE | t statistics | p value |
|  | **anterior MTG** |  |  |  |  |  |  |  |  |  |  |  |
| Intercept | 0.08 | 0.29 | 0.29 | 0.776 | -0.09 | 0.35 | -0.24 | 0.811 | 0.21 | 0.32 | 0.66 | 0.519 |
| Gender | -0.38 | 0.89 | -0.43 | 0.677 | 0.39 | 1.08 | 0.36 | 0.727 | -0.96 | 0.99 | -0.97 | 0.348 |
| Age | 0.41 | 0.36 | 1.14 | 0.274 | 0.17 | 0.44 | 0.38 | 0.712 | -0.07 | 0.40 | -0.18 | 0.862 |
| RT | **-0.67** | **0.27** | **-2.49** | **0.026** | 0.00 | 0.33 | -0.01 | 0.993 | 0.08 | 0.30 | 0.26 | 0.799 |
|  | **posterior MTG** |  |  |  |  |  |  |  |  |  |  |  |
| Intercept | 0.16 | 0.30 | 0.54 | 0.596 | -0.02 | 0.35 | -0.07 | 0.946 | 0.14 | 0.33 | 0.42 | 0.681 |
| Gender | -0.74 | 0.93 | -0.80 | 0.439 | 0.11 | 1.06 | 0.10 | 0.921 | -0.62 | 1.00 | -0.62 | 0.548 |
| Age | 0.05 | 0.38 | 0.13 | 0.902 | 0.13 | 0.43 | 0.31 | 0.764 | -0.06 | 0.41 | -0.15 | 0.885 |
| RT | **-0.65** | **0.28** | **-2.30** | **0.038** | -0.25 | 0.32 | -0.78 | 0.448 | 0.23 | 0.30 | 0.75 | 0.463 |
|  | **anterior temporal clusters in color anomia** |  |  |  |  |  |  |  |  |  |  |  |
| Intercept | 0.35 | 0.26 | 1.32 | 0.209 | 0.22 | 0.33 | 0.67 | 0.516 | 0.07 | 0.32 | 0.20 | 0.841 |
| Gender | -1.57 | 0.81 | -1.93 | 0.074 | -1.00 | 1.03 | -0.98 | 0.345 | -0.30 | 1.00 | -0.30 | 0.769 |
| Age | 0.07 | 0.33 | 0.21 | 0.841 | -0.17 | 0.42 | -0.42 | 0.684 | 0.26 | 0.41 | 0.65 | 0.526 |
| RT | **-0.74** | **0.25** | **-3.02** | **0.009** | -0.36 | 0.31 | -1.14 | 0.272 | 0.05 | 0.30 | 0.16 | 0.879 |

## Notes

### Competing Interest Statement

The authors have declared no competing interest.

### Summary of Updates

-Correction of spelling errors -Replacing previous figures with lower quality equivalents for easier transfers

